# Systematic Profiling of Temperature- and Retinal-Sensitive Rhodopsin Variants by Deep Mutational Scanning

**DOI:** 10.1101/2021.03.26.437229

**Authors:** Andrew G. McKee, Charles P. Kuntz, Joseph T. Ortega, Hope Woods, Francis J. Roushar, Jens Meiler, Beata Jastrzebska, Jonathan P. Schlebach

**Author notes:** Authors contributed equally (Andrew G. McKee, Charles P. Kuntz).

## Abstract

Membrane protein variants with diminished conformational stability often exhibit enhanced cellular expression at reduced growth temperatures. The expression of “temperature-sensitive” variants is also typically sensitive to corrector molecules that bind and stabilize the native conformation. In this work, we employ deep mutational scanning to compare the effects of reduced growth temperature and an investigational corrector (9-*cis*-retinal) on the plasma membrane expression of 700 rhodopsin variants in HEK293T cells. We find that the change in expression at reduced growth temperatures is correlated with the response to retinal among variants bearing mutations within a hydrophobic transmembrane domain (TM2). The most sensitive variants within this helix appear to disrupt a network of hydrogen bonds that stabilizes a native helical kink. By comparison, mutants that alter a polar transmembrane domain (TM7) exhibit weaker responses to temperature and retinal that are poorly correlated. Statistical analyses suggest this insensitivity primarily arises from an abundance of mutations that enhance its membrane integration, stabilize its native conformation, and/ or perturb the retinal binding pocket. Finally, we show that the characteristics of purified temperature- and retinal-sensitive variants suggest that the proteostatic effects of retinal may be manifested during translation and cotranslational folding. Together, our findings elucidate various factors that mediate the sensitivity of genetic variants to temperature and to small molecule correctors.

## Introduction

Eukaryotic membrane proteins are prone to misfolding, and mutations that enhance their propensity to misfold are associated with a myriad of diseases of aberrant protein homeostasis.^1^ These destabilized variants are typically recognized and degraded by the ER associated degradation (ERAD) pathway, which ultimately reduces the expression of functional protein at the plasma membrane.^2^ Efforts to identify and characterize these misfolded, disease-linked variants have revealed that many partially recover their expression when cells are incubated below their physiological growth temperature (37°C).^3–7^ Expressing proteins at lower temperatures is believed to compensate for the effects of these mutations by increasing the thermodynamic preference for the native conformation, which is typically maximized near room temperature.^8^ This temperature-sensitive phenotype has long served as a marker for destabilizing mutations that induce misfolding and degradation (class II). Nevertheless, class II mutations within integral membrane proteins can cause a spectrum of conformational defects,^9^ and it is currently unclear which types can be reversed by temperature or other factors that influence proteostasis.

The expression of many tempterature-sentive variants can also be restored by small molecule correctors that preferentially bind and stabilize the native conformation,^10^ some of which are in current use for the treatment of cystic fibrosis.^11–13^ However, response to these compounds varies considerably among class II variants for reasons that remain unclear as a result of the heterogeneity of their molecular defects^14^ and the challenges associated with membrane protein folding measurements.^15^ To gain insights into the basis of these variations, we recently surveyed mutagenic trends in the clas A G-protein coupled receptor rhodopsin, the misfolding of which causes retinitis pigmentosa (RP).^16^ With the use of deep mutational scanning (DMS),^17^ we analyzed the proteostatic effects of 808 rhodopsin mutations and how they respond to the investigational corrector 9-*cis*-retinal-^18–20^ a photostable analog of rhodopsin’s native cofactor (11-*cis*-retinal). These compounds bind to the folded opsin apoprotein with high affinity^21^ and form a covalent Schiff base with a conserved lysine (K296). Our investigations have collectively revealed that the plasma membrane expression (PME) of rhodopsin is particularly sensitive to mutations within its seventh transmembrane domain (TM7),^20^ which is intrinsically prone to cotranslational misfolding.^19^ Moreover, variants bearing mutations within TM7 appear to be less sensitive to the proteostatic effects of retinal relative to mutations within its second transmembrane domain (TM2), which is considerably more hydrophobic.^20^ These observations suggest mutations within different regions of the molecule may have divergent pharmacological profiles as a result of their distinct conformational effects. Nevertheless, it remains challenging to delineate the effects of mutations on cotranslational folding, post-translaitional folding, and the intrinsic binding energetics.

In this work, we utilize DMS to survey the temperature-sensitivity of 700 combined TM2 and TM7 variants in relation to their response to 9-*cis*-retinal. Using a panel of computational and structural tools, we identify various classes of temperature- and/ or retinal-sensitive mutations. Consistent with expectations, we find a robust statistical correlation between the degree of temperature- and retinal-sensitivity amoung TM2 variants. Moreover, we identify a cluster of mutations within this helix that render rhodopsin both temperature- and retinal-sensitive by disrupting a native helical kink. An analysis of the crystal structures of G90D and T94I rhodopsin,^22,23^ which cause congenital stationary night blindness (CSNB), confirms that polar mutations within this region disrupt the native hydrogen bonds that stabilize this kink. By comparison, we find fewer temperature-sensitve mutations within TM7, and show that temperature-sensitivity does not coincide with retinal-sensitivity within this region. Statistical trends within these data suggest this disconnect arises from the differential effects of these mutations on the stability of the native conformation, the fidelity of cotranslational folding, and/ or the integrity of the retinal binding pocket. Together, these results provide insights into how the molecular defects caused by different classes of mutations ultimately influence their sensitvity to temperature and to correctors. Moreover, this unbiased survey of missense variants provides a molecular context to interpret the proteostatic effects and pharmacological profiles of the mutations that are known to cause RP and CSNB.

## Results

### Identification of Temperature-Sensitive Opsin Variants

To identify and characterize temperature-sensitive rhodopsin variants, we measured changes in the PME of several hundred variants at reduced growth temperature by DMS. We first utilized a genetically modified HEK293T cell line bearing a single genomic recombination site^24^ to produce a pool of stable cells that collectively express a mixed library of TM2 variants, a mixed library of TM7 variants, or WT rhodopsin only. The surface immunostaining of stable cells expressing WT opsin increased by an average of 31% when the growth temperature was reduced to 27°C for 24 hours (Fig. 1A). Reducing the growth temperature also increased the average surface immunostaining intensity of recombinant cells expressing either the collection of TM2 variants (+13%, Fig. 1B) or TM7 variants (+8%, Fig. 1C) contained within each genetic library. Cells expressing these rhodopsin variants exhibit a relatively modest change in average intensity at 27°C as a result of a minor subpopulation that expresses insensitive variants that exhibit minimal surface immunostaining under either condition (Fig. 1 B & C). Nevertheless, the relative proportion of these cells decreases at reduced temperature (Fig. 1 B & C), which suggests many of these (likely class II) variants regain expression at 27°C. These observations suggest that, like WT, many variants bearing mutations within these TM domains exhibit enhanced PME when the growth temperature is reduced to 27°C.

**Figure 1.**
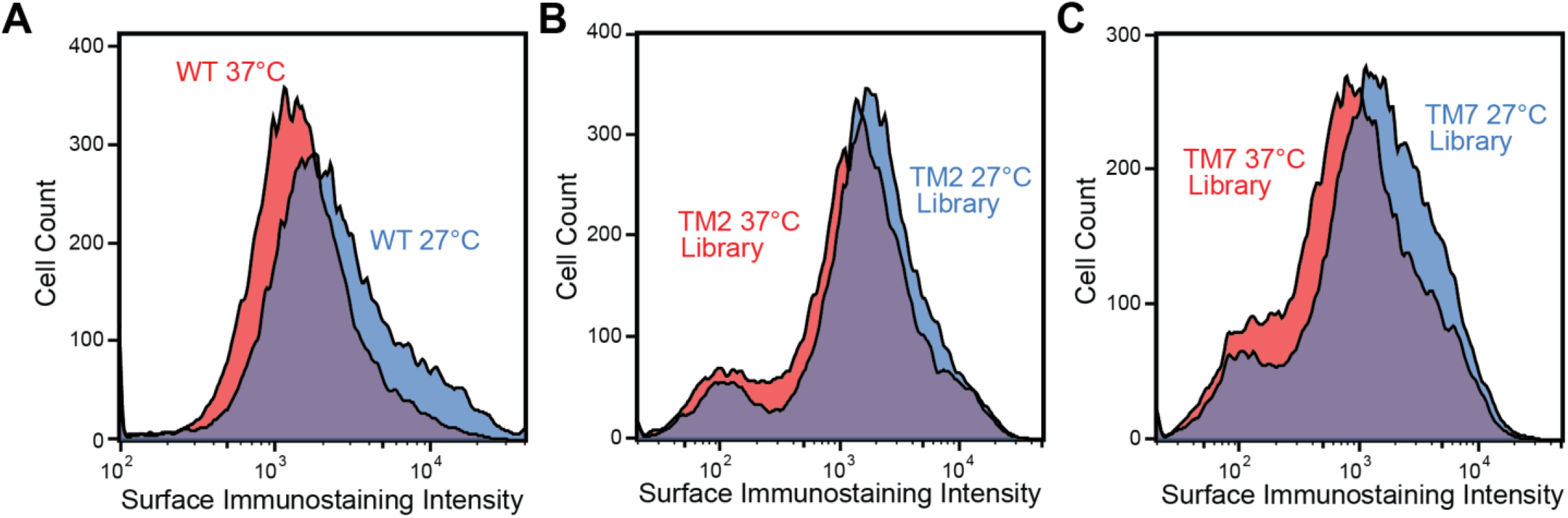
Impact of Growth Temperature on the Cell Surface Immunostaining of Opsin. Histograms depict the distribution of opsin surface immunostaining intensities among recombinant HEK293T cells expressing A) WT rhodopsin, B) individual TM2 variants, or C) individual TM7 variants grown at 37°C (red) or 27°C (blue). Traces were created from a representative set of 25,000 cellular intensities.

To identify specific variants that exhibit enhanced expression at 27°C, we first utilized fluorescence activated cell sorting (FACS) to fractionate the cellular libraries grown at each temperature according to the distribution of surface immunostaining intensities. We then used deep sequencing to track the expressed variants within each fraction and to estimate each of their surface immunostaining intensities as was previously described.^20^ To facilitate the comparison of these intensity values across replicates and conditions, we previously opted to normalize the value for each variant by that of WT.^20^ However, the temperature-sensitive expression of WT opsin undermines its utility as an internal control (Fig. 1A). As an alternative, we used the average intensity values for the collection of truncated nonsense variants within each library to normalize the intensity values of each missense variant. Heatmaps depicting normalized intensity values at 27°C relative to those at 37°C reveal which variants exhibit the largest increase in PME at reduced growth temperature (Fig. 2 A & B). These data reveal few generalizable trends about the sensitivity of TM2 or TM7 variants, which suggests temperature-sensitivity depends on the structural context of their mutations.

**Figure 2.**
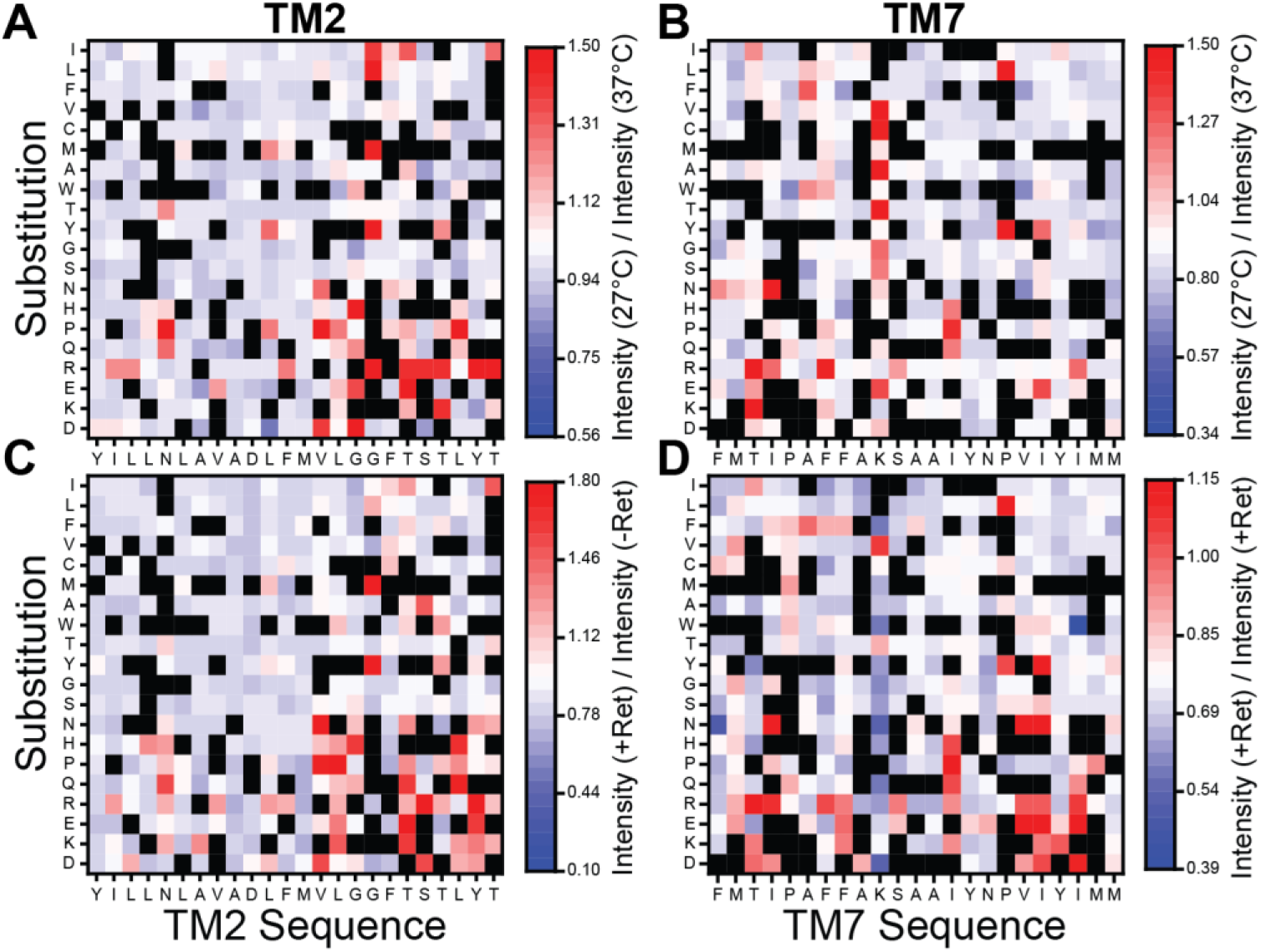
Temperature- and Retinal-Sensitivity of Rhodopsin Variants. Deep mutational scanning was used to measure the change in the plasma membrane expression of rhodopsin variants at reduced growth temperature or in the presence of 9-*cis*-retinal. A) A heatmap depicts the surface immunostaining intensities for a collection of TM2 variants bearing individual amino acid substitutions (y-coordinate) at each residue (x-coordinate) at 27°C normalized relative to the corresponding values at 37°C. B) A heatmap depicts the surface immunostaining intensities for TM7 variants at 27°C normalized relative to the corresponding values at 37°C. C) A heatmap depicts the surface immunostaining intensities for TM2 variants in the presence of 5 μM 9-*cis*-retinal at 37°C normalized relative to the corresponding values in the absence of 9-*cis*-retinal at 37°C. D) A heatmap depicts the surface immunostaining intensities for TM7 variants in the presence of 5 μM 9-*cis*-retinal at 37°C normalized relative to the corresponding values in the absence of 9-*cis*-retinal at 37°C. Values represent the average from two biological replicates and color bars indicate the scale of the observed effects under each condition. Red indicates an increase in PME, blue indicates a reduction in PME, and white indicates no change in PME under each condition. Black squares indicate a lack of reliable data.

Temperature-sensitivity is typically associated with unstable variants that are poorly expressed, and it is generally assumed that their enhanced expression arises from an increase in thermodynamic stability and a corresponding increase in the fraction of folded protein at reduced growth temperatures. In this case, variants bearing destabilizing mutations that decrease PME should exhibit the largest increase in expression at 27°C. Indeed, a plot of intensity ratios against the relative PME of each variant at 37°C reveals that temperature-sensitivity trends upwards as expression levels decrease (Fig. 3A). However, this uptick in expression is only statistically significant among TM2 variants with diminished expression, and there appear to be relatively few temperature-sensitive TM7 variants overall (Fig. 3B). Together, these results reveal trends in the temperature-sensitivity of 700 opsin variants, and hint at underlying differences in the molecular effects of mutations in TM2 and TM7.

**Figure 3.**
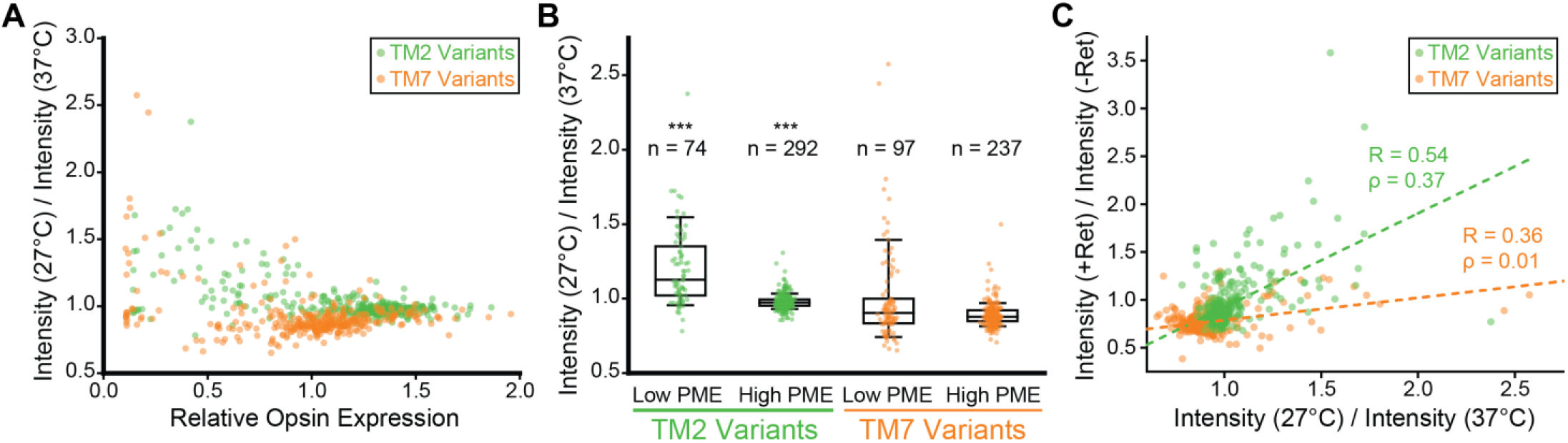
Temperature-Sensitivity of Opsin Variants in Relation to Plasma Membrane Expression and Retinal Response. The temperature sensitivity of opsin variants is examined in relation to their relative plasma membrane expression levels at 37°C and their corresponding response to 9-*cis*-retinal. A) The ratio of the surface immunostaining intensity at 27°C relative to that at 37°C is plotted against the corresponding surface immunostaining values normalized relative to WT at 37°C for TM2 (green) and TM7 (orange) variants. B) A box and whisker plot depicts the distribution of intensity ratios among TM2 (green) and TM7 (orange) variants that were classified as either high or low PME variants based on a relative opsin expression cutoff value of 0.9 (see ref 17). Whiskers reflect the 10^th^ and 90^th^ percentile values, and the edges of the boxes reflect the 25^th^ and 75^th^ percentile values. The horizontal line within the box reflects the median intensity ratio. The distributions of intensity ratios among high and low PME variants of TM2 (***) were found to be statistically distinct at a confidence level of 0.001 by to a two-tailed Mann-Whitney U test. C) The ratio of the surface immunostaining intensity in the presence of 5 μM retinal at 37°C relative to that in the absence of retinal at 37°C is plotted against the corresponding ratio of the surface immunostaining intensity at 27°C relative to that at 37°C for TM2 (green) and TM7 (orange) variants. A linear fit of the intensity ratio data for TM2 variants (green dashes, Pearson’s R = 0.54) and TM7 variants (orange dashes, Pearson’s R = 0.36) are shown for reference.

### Relationships between Temperature- and Retinal-Sensitivity

To evaluate the coincidence between sensitivity to temperature and to correctors, we utilized our previous measurements^20^ to calculate the change in the surface immunostaining of these variants in the presence of 5 μM 9-*cis*-retinal. Heatmaps depicting the change in the immunostaining intensity for each variant reveal few generalizable trends about retinal-sensitive mutations (Fig. 2 C & D), which suggests this property also depends on structural context.

Mutations that introduce polar side chains within the C-terminal residues of TM2 generally render rhodopsin sensitive to both temperature and retinal (Fig. 2 A & C). These residues are directly adjacent to a rigid region within the rhodopsin structure that makes key contributions to its stability.^25^ However, a simulated thermal denaturation^26,27^ of the native rhodopsin ensemble suggests the C-terminal residues are actually more dynamic than the N-terminal residues of TM2 (Fig. S1). Interestingly, these dynamic residues surround a native kink that is mediated by two consecutive glycine residues (G89 & G90, Fig. 2 A & C, Fig. 4 A & B). This kink is stabilized by a network of hydrogen bonds established between G89 (backbone C=O), G90 (backbone C=O), S93 (backbone NH, sidechain OH), and T94 (backbone NH & sidechain OH, Fig. 4C). Notably, there are several pathogenic mutations that either introduce a polar side chain or remove a native polar side chain near this kink. Three of these mutations cause RP (V87D, G89R, and T89I) and result in a loss of PME.^20^ Two others that cause CSNB (G90D & T94I) feature non-native tertiary contacts that increase expression yet disrupt retinal binding and/ or the native photocycle. Nevertheless, crystal structures of these two variants show that, while the helical kink is maintained by surrounding tertiary contacts, the hydrogen bonding network that locally stabilizes it is disrupted (Fig. 4 D & E).^22,23^ Based on these observations, we suspect most polar mutations within this region render the protein temperature- and retinal-sensitive by disrupting hydrogen bonding and destabilizing this native kink (see *Discussion*).

**Figure 4.**
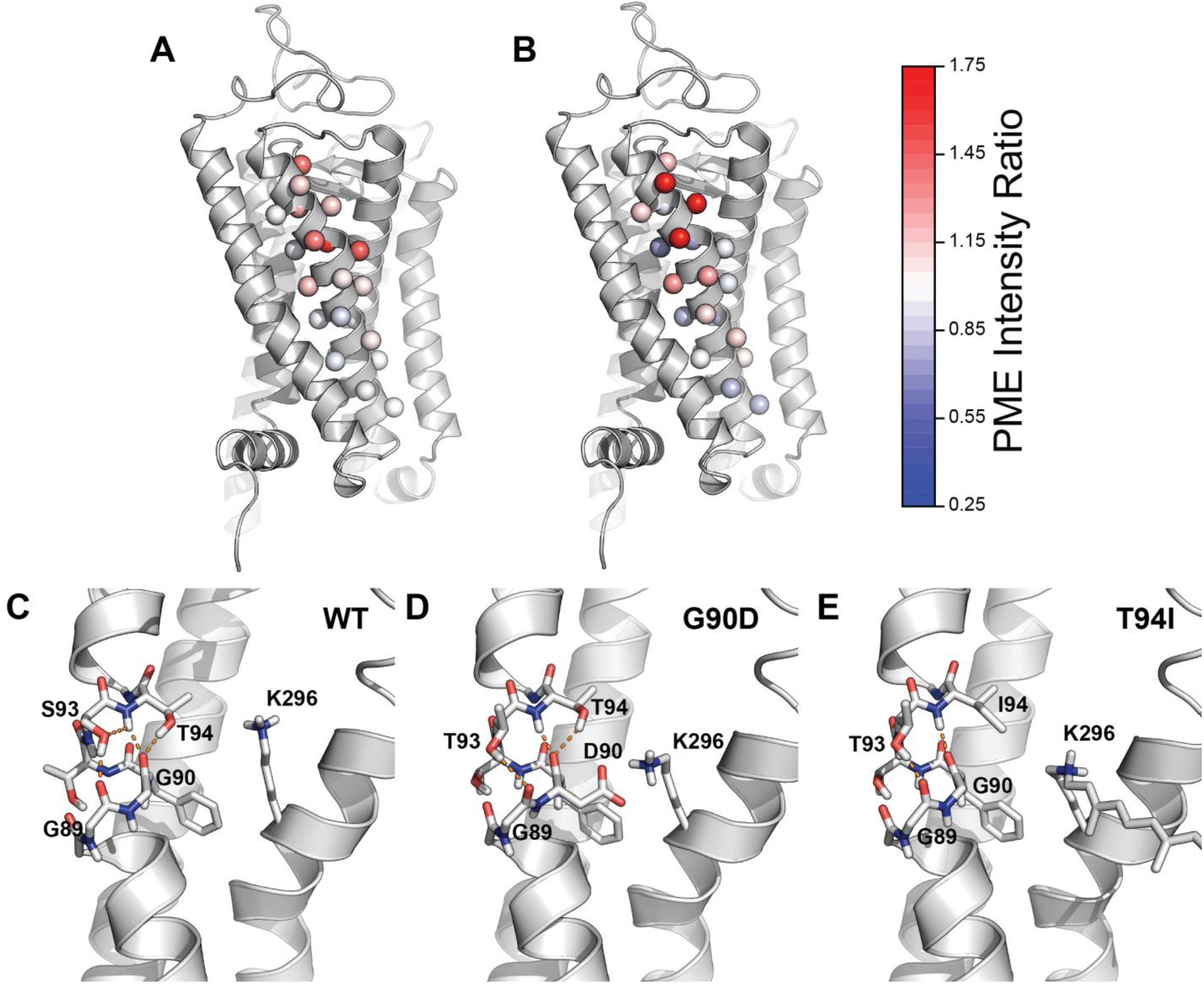
Structural Context of Temperature- and Retinal-Sensitive Mutations within TM2. Temperature- and retinal-sensitivity measurements for charged variants within TM2 are projected onto an active state crystal structure of rhodopsin (PDB 5W0P). A) The ratio of the surface immunostaining intensity at 27°C relative to that at 37°C was averaged across charged substitutions in TM2 and projected as colored spheres on to the Cβ (or glycine hydrogen). The magnitude of the intensity ratios is shown in the color bar. B) The ratio of the surface immunostaining intensity in the presence of 5 μM 9-*cis*-retinal relative to that in the absence of retinal was averaged across charged substitutions in TM2 and projected as colored spheres on to the Cβ (or glycine hydrogen). The magnitude of the intensity ratios is shown in the color bar. C) The hydrogen bond network stabilizing the kink in human WT TM2 is shown (PDB 5W0P). D) The hydrogen bond network stabilizing the kink in TM2 of bovine G90D rhodopsin is shown (PDB 4BEY). E) The hydrogen bond network stabilizing the kink in TM2 of bovine T94I rhodopsin is shown (PDB 5EN0). Hydrogen bonds are connected by orange dashes. Bovine rhodopsin (94% identical to human) contains a threonine instead of a serine at residue 93.

Across all TM2 variants, the change in expression in the presence of 9-*cis*-retinal is statistically correlated with the change in expression at 27°C (Fig. 3C, Pearson’s R = 0.54, *m* = 0.98 ± 0.08). In contrast, the response of TM7 variants to retinal appears to diverge from their response to a reduction in growth temperature (Fig. 2 B & D). Moreover, the correlation between the intensity ratios of TM7 variants is considerably weaker, and increases in surface immunostaining at 27°C generally coincides with a smaller change in the presence of retinal (Fig. 3C, Pearson’s R = 0.36, *m* = 0.23 ± 0.03). We note that, if changes in PME reflect the impacts of mutations on the free energy of folding and the corresponding fraction of folded protein, then ideal trends should be nonlinear. We therefore calculated Spearman’s rank correlation coefficients, which do not assume linearity, in order to evaluate the statistical significance of the coincidence between temperature- and retinal-sensitivity among these variants. Rank correlation coefficients suggest the coincidence between temperature- and retinal-sensitivity is highly significant among TM2 variants (ρ = 0.37. p = 3.6×10^−13^) but not among TM7 variants (ρ = 0.01, p = 0.79). Together, these observations reveal that temperature-sensitivity only coincides with retinal-sensitivity among mutations within certain regions of the protein.

### Structural and Energetic Basis for the Coupling between Temperature- and Retinal-Sensitivity

We suspect the decoupling of temperature-sensitivity from retinal-sensitivity among TM7 variants arises from a distinction in the underlying conformational effects of these mutations. In the following section, we analyze deviations in the expression profiles of certain sub-sets of variants in order to test a series of hypotheses related to the origins of temperature- and retinal-sensitivity.

We first hypothesized that certain mutations in TM7 render rhodopsin expression insensitive to changes in temperature and/ or retinal by enhancing the stability of the native conformation and increasing the fraction of folded apoprotein. To test this hypothesis, we analyzed mutant sensitivity profiles in relation to their predicted impacts on conformational stability. Both technical challenges^15^ and the sheer volume of data preclude experimental approaches to measure the stability of the 700 rhodopsin mutants characterized herein. As an alternative, we calculated Rosetta ΔΔG values^28^ to estimate the effects of each of these mutations on the free energy of the native conformation.^20^ Despite quantitative limitations,^29^ ensembles of stability predictions remain useful for the identification of energetic trends within large data sets.^20,30^ Using these ΔΔG values, we applied a series of increasingly stringent cutoffs to filter out neutral or stabilizing mutations and define subsets of TM7 variants that are enriched with destabilizing mutations. For both TM2 and TM7 variants, the rank correlation between temperature- and retinal-sensitivity improves as the stringency of the cutoff value increases (Table 1). Moreover, this correlation is statistically significant among the subset of TM7 variants predicted to increase the free energy of folding by more than 3.0 Rosetta energy units (REU, Table 1). Consistent with expectations, median intensity ratios associated with the response to both temperature and retinal are higher among destabilized variants (ΔΔG > 5.0 REU) in both TMs 2 & 7 (Fig. 5). Together, these findings suggest the insensitivity of TM7 variants arises, in part, from the fact that the native rhodopsin fold appears to be stabilized by a relatively large proportion of random mutations within this domain.

**Table 1.** Coincidence Between Temperature and Retinal Response Among Variant Sub-Classes.

| Parsing | TM2 Variants |  |  | TM7 Variants |  |  |
| --- | --- | --- | --- | --- | --- | --- |
| | Number of Variants | Spearman's $\rho$ | p- value | Number of Variants | Spearman's $\rho$ | p- value |
| All Variants | 366 | 0.37 | $3.6 \times 10^{-13}$ | 334 | 0.01 | 0.79 |
| Predicted Change in the Free Energy of Folding (Rosetta $\Delta\Delta G$) | | | | | | |
| > +1.0 REU | 240 | 0.39 | $2.2 \times 10^{-10}$ | 167 | 0.09 | 0.23 |
| > +3.0 REU | 160 | 0.44 | $6.9 \times 10^{-9}$ | 91 | 0.20 | 0.05 |
| > +5.0 REU | 90 | 0.54 | $4.3 \times 10^{-8}$ | 56 | 0.43 | $1.0 \times 10^{-3}$ |
| Predicted Change in the Free Energy of the Membrane Integration (Biological Hydrophobicity Scale $\Delta\Delta G$) | | | | | | |
| > +0.50 kcal/ mol | 149 | 0.38 | $1.4 \times 10^{-7}$ | 109 | 0.17 | 0.08 |
| > +0.75 kcal/ mol | 109 | 0.44 | $1.7 \times 10^{-6}$ | 77 | 0.14 | 0.21 |
| > +1.0 kcal/ mol | 69 | 0.42 | $2.9 \times 10^{-4}$ | 48 | 0.32 | 0.03 |
| Distance from C$_{\alpha}$ to Retinal Binding Pocket* | | | | | | |
| >10 Å | 366 | 0.37 | $3.6 \times 10^{-13}$ | 285 | 0.11 | 0.07 |
| >12 Å | 366 | 0.37 | $3.6 \times 10^{-13}$ | 208 | 0.19 | $5.2 \times 10^{-3}$ |
| >14 Å | 301 | 0.40 | $8.0 \times 10^{-13}$ | 166 | 0.22 | $5.4 \times 10^{-3}$ |

\*Distances were calculated based on the crystal structure of bovine rhodopsin (PDB 3C9L). The position of the binding pocket was defined as the centroid of the retinal cofactor.

**Figure 5.**
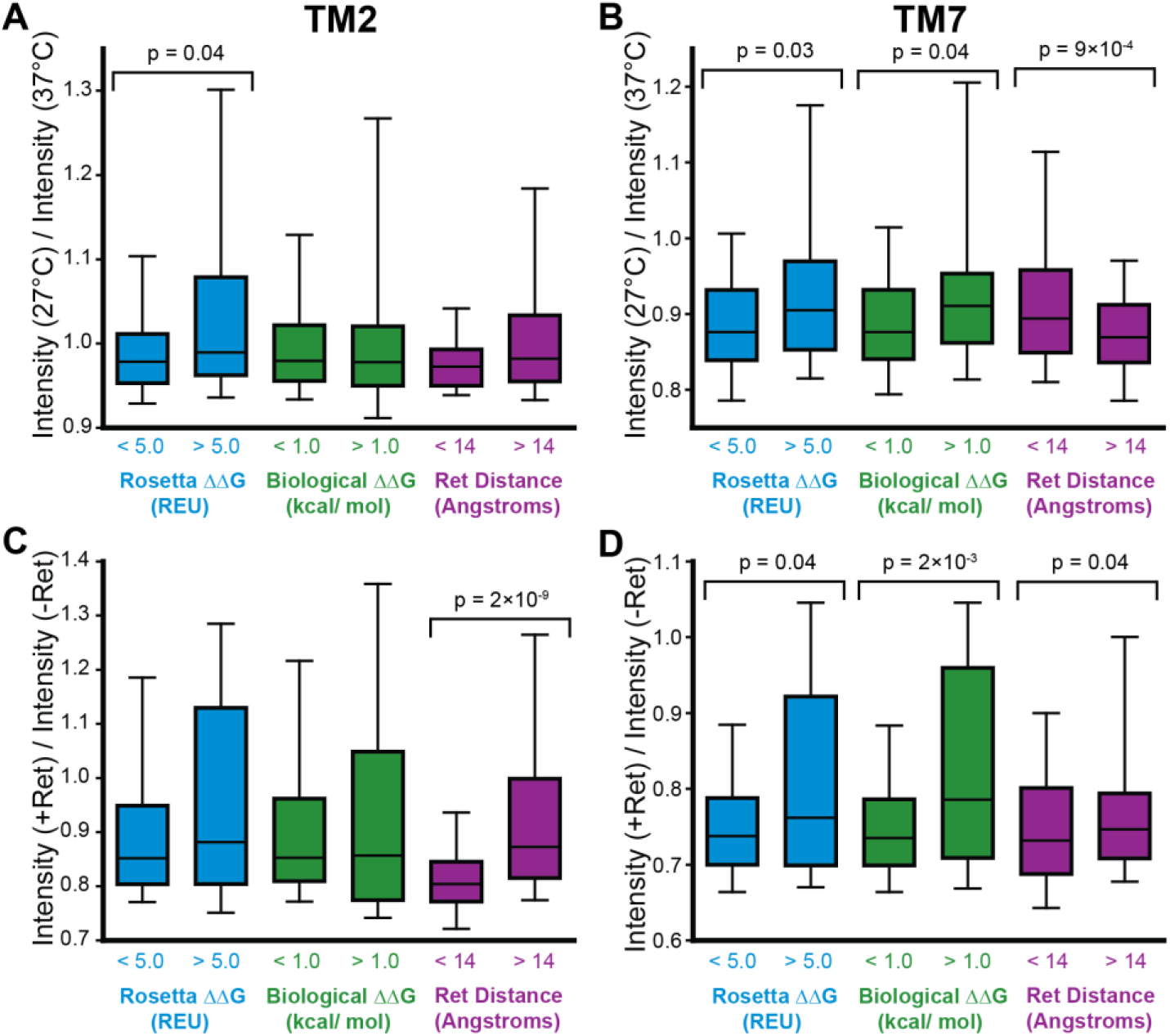
Sensitivity of Mutations that Disrupt Folding, Membrane Integration, or the Retinal Binding Pocket. The distribution of the ratio of the surface immunostaining intensity at 27°C relative to that at 37°C among TM2 (A) and TM7 (B) variants that were classified based on Rosetta ∆∆G values (blue), biological hydrophobicity scale ΔΔG values (green), or distance from the retinal binding pocket (purple) are depicted using a box and whisker plot. The distribution of the ratio of the surface immunostaining intensity in the presence of 5 μM 9-*cis*-retinal at 37°C relative to that in the absence of retinal at 37°C among TM2 (C) and TM7 (D) variants that were classified based on Rosetta ΔΔG values (blue), biological hydrophobicity scale ΔΔG values (green), or distance from the retinal binding pocket (purple) are depicted using a box and whisker plot. Whiskers reflect the 10^th^ and 90^th^ percentile values, and the edges of the boxes reflect the 25^th^ and 75^th^ percentile values. The horizontal line within the box reflects the median intensity ratio. A two-tailed Mann-Whitney U-test was used to compare the distribution of intensity ratios among variants above and below each cutoff, and p-value are shown for each case in which the differences in the distributions were found to be statistically significant.

We previously found that TM7 is topologically frustrated, and showed that rhodopsin expression is highly sensitive to mutations that alter its membrane integration efficiency.^19^ We therefore hypothesized that this class of mutations could also potentially render rhodopsin expression insensitive to further stabilization by temperature or retinal. To test this hypothesis, we utilized the biological hydrophobicity scale^31^ to filter out mutations that are predicted to enhance the efficiency of the translocon-mediated membrane integration of TM7. Regardless of the applied cutoff, the rank correlation between temperature- and retinal-sensitivity is stronger among mutations in TM7 that are predicted to disrupt its translocon-mediated membrane integration (Table 1). Though narrowing the pool of mutants reduces statistical power, the strength and significance of the correlation between these variables is most pronounced among the most highly destabilized variants (ΔΔG > 1.0 kcal/ mol, ρ = 0.32, p = 0.03). Moreover, this sub-set of variants exhibits an enhanced response to both temperature and retinal (Fig. 5 B & D). However, a fitted trend line suggests the response to retinal among topologically destabilized TM7 variants (*m* = 0.5 ± 0.1) is smaller in magnitude relative to that of TM2 variants (*m* = 0.98 ± 0.08). For reference, the correlation between temperature- and retinal-sensitivity is also slightly stronger among this class of mutations within TM2 (Table 1), though this sub-set of variants do not exhibit greater sensitivity to temperature or retinal (Fig. 5 A & C). It should be noted that only 35% of the TM7 variants with biological hydrophobicity ΔΔG values > 1.0 kcal/ mol also have a Rosetta ΔΔG value > 5.0 REU. This modest overlap suggests the impacts of mutations on topology and stability are largely independent. Together, these observations suggest the expression of variants bearing mutations that enhance the membrane integration of TM7 are typically less temperature- and/ or retinal-sensitive.

TM7 lines the retinal binding pocket and contains K296, which forms a functionally essential Schiff base with the retinal cofactor. We therefore hypothesized that certain TM7 mutations dampen the response to retinal by compromising this binding pocket. Estimating the impacts of mutations on binding affinity is challenging. Nevertheless, we presume that mutations near the retinal pocket are more likely to perturb binding- a classic assumption in pharmacology. We therefore used the proximity of native residues to retinal as a metric to filter out mutations that are most likely to perturb binding. We calculated the distance from the α carbon of each residue to the centroid position of retinal using a high-resolution structure of bovine rhodopsin (PDB 3C9L). We then applied a series of distance-based cutoffs to analyze the correlation between temperature- and retinal-sensitivity among TM7 variants that are increasingly distant from the retinal binding pocket. The correlation between the response of TM7 variants to temperature and retinal is more pronounced among variants that are distant from the retinal binding pocket (Table 1). Moreover, these rank correlations become highly significant among variants that are located > 12Å away from retinal (Table 1). Mutations distant from the binding pocket exhibit an enhanced response to retinal (Fig. 5D), but a weaker response to temperature (Fig. 5B). We attribute this later observation to the fact that mutations are more distant from the protein core and are therefore less likely to introduce packing defects that could compromise stability. For reference, residues within TM2 are further away from the retinal binding pocket, on average, relative to those in TM7, and far fewer TM2 mutations are removed by these criteria (Table 1). Nevertheless, the 65 TM2 mutations within 14 Å of the retinal binding pocket exhibit a diminished response to retinal (Fig. 5C), and removing them slightly increases the strength of the correlation between temperature- and retinal sensitivity (Table 1). These observations suggest the direct impacts of mutations on retinal binding also contributes to deviations between temperature- and retinal-sensitivity.

### Retinal Binding and Thermal Stability of Purified Rhodopsin Variants

Our analyses collectively suggest temperature- and retinal-sensitivity stems from variations in the fidelity of both cotranslational and post-translational folding. To evaluate the relative contribution of these co- and post-translational effects, we characterized the purified, full-length forms of three TM2 variants (A80Q, G90M, and Y96S) and three TM7 variants (F287N, T289I, and N302Y) that vary with respect to their sensitivity to temperature and retinal (Table 2). Among these variants, we find that G90M and T289I exhibit particularly robust increases in PME in the presence of retinal and at 27°C (Figs. 6 A & B, Table 2). To evaluate the propensity of full-length variants to bind retinal and form the native structure we expressed each protein in HEK293T cells, harvested cellular membranes, treated membranes with 9-*cis*-retinal, then purified each regenerated rhodopsin pigment. Comparison of full-length variant spectra and the ratio of their absorbance at 280 nm to 485 nm (a measure of regeneration efficiency) suggests that both G90M and T289I compromise post-translational folding and/ or binding (Fig. 6C, Table 2). By comparison, the regeneration of G90M and T289I rhodopsins was less efficient relative to that of variants with expression profiles that are more comparable to WT (Table 2). These observations suggest retinal sensitivity does not track with the post-translational effects of these mutations on retinal binding and/ or folding. Interestingly, formation of the native pigment is more efficient when retinal is added to live cells actively expressing G90M and T289I rhodopsin prior to purification (Fig. 6D). This observation suggests the proteostatic effects of retinal are derived from interactions that occur during translation and cotranslational folding. It should also be noted that purified G90M and T289I pigments exhibit diminished kinetic stability at 27°C relative to WT (Fig. S2). Co-translational regeneration of these pigments increases their kinetic stability, though mutant pigments are still less stable than WT (Fig. S2). This observed instability of the regenerated pigments suggests the temperature-sensitivity of these variants is unlikely to arise from an enhanced conformational stability and/ or binding affinity in these full-length variants.

**Table 2.** Cellular and Biochemical Properties of Transiently Expressed Rhodopsin Variants.

| Variant | Relative PME<br>at 37°C* | Relative PME<br>at 37°C + 5 μM Retinal** | Relative PME<br>at 27°C*** | Regeneration Efficiency<br>(A<sub>280</sub> / A<sub>485</sub>) |
| --- | --- | --- | --- | --- |
| WT | 1.0 | 1.4 ± 0.2 | 1.7 ± 0.1 | 2.97 |
| A80Q | 0.5 ± 0.1 | 1.8 ± 0.2 | 2.1 ± 0.2 | 5.18 |
| G90M | 0.5 ± 0.09 | 3.9 ± 0.4 | 3.6 ± 0.6 | 6.90 |
| Y96S | 0.6 ± 0.04 | 1.9 ± 0.2 | 1.7 ± 0.09 | 2.32 |
| F287N | 0.7 ± 0.06 | 1.5 ± 0.1 | 1.9 ± 0.2 | 2.50 |
| T289I | 0.6 ± 0.04 | 2.9 ± 0.8 | 4.2 ± 1.0 | 12.4 |
| N302Y | 0.9 ± 0.1 | 1.2 ± 0.09 | 1.8 ± 0.04 | 3.99 |

\*Mean surface immunostaining intensities of HEK293 cells transiently expressing variants were normalized relative to those of cells expressing WT rhodopsin. Values represent the average of three biological replicates ± the standard deviation.
\*\*Mean surface immunostaining intensities of HEK293 cells transiently expressing variants in the presence of 5 μM 9-*cis*-retinal were normalized relative to the variant intensity in the absence of retinal. Values represent the average of three biological replicates ± the standard deviation.
\*\*\*Mean surface immunostaining intensities of HEK293 cells transiently expressing variants at 27°C were normalized relative to the value for the variant intensity at 37°C. Values represent the average of three biological replicates ± the standard deviation.

**Figure 6.**
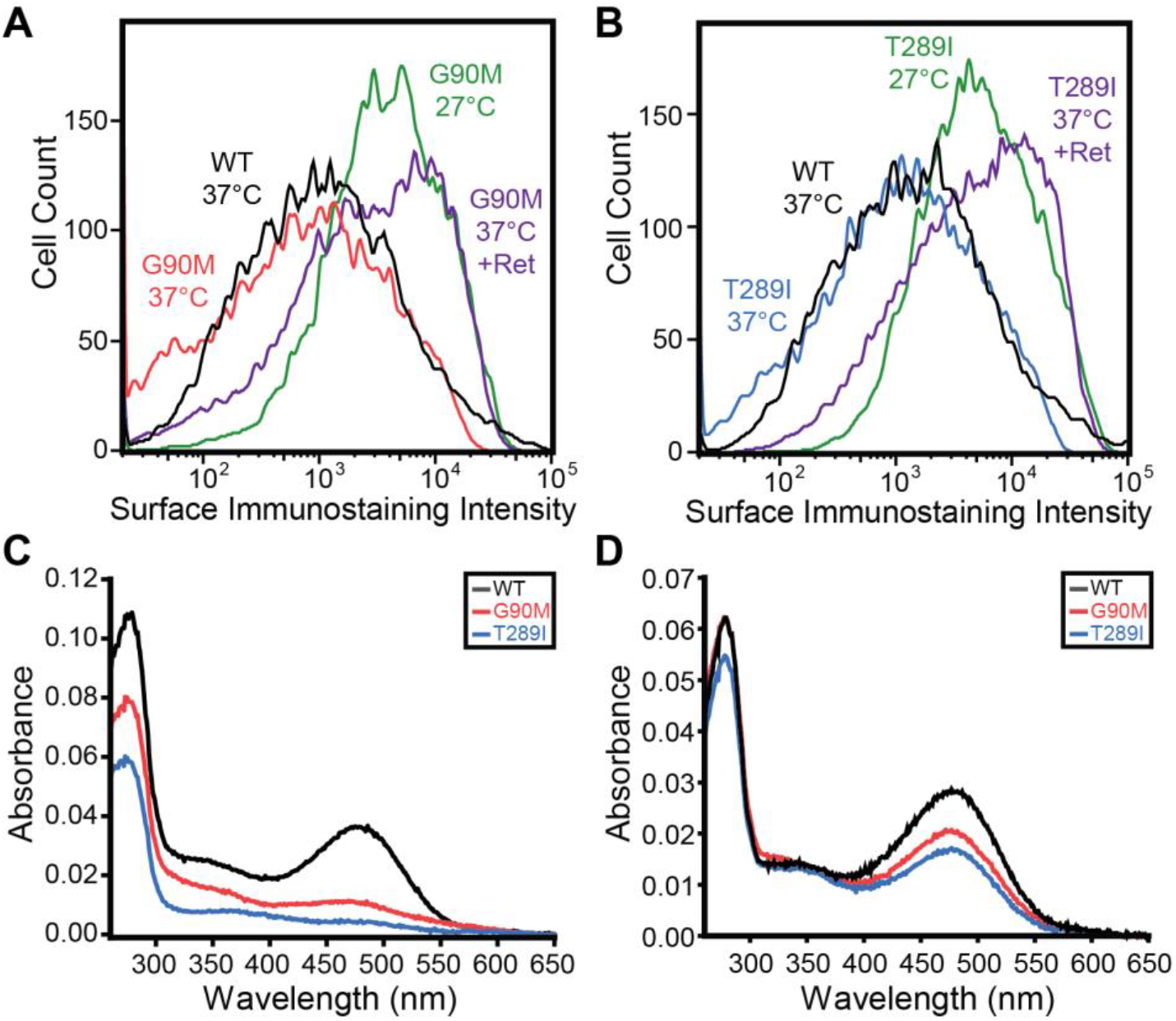
Expression and Regeneration of G90M and T289I Rhodopsins. A) A histogram depicts flow cytometry measurements of the surface immunostaining intensity of HEK293T cells transiently expressing G90M rhodopsin at 37°C (red), 27°C (green), or at 37°C in the presence of 5μM 9-*cis*-retinal (purple). The surface immunostaining intensities of cells transiently expressing WT rhodopsin at 37°C (black) is shown for reference. B) A histogram depicts flow cytometry measurements of the surface immunostaining intensity of HEK293T cells transiently expressing T289I rhodopsin at 37°C (blue), 27°C (green), or at 37°C in the presence of 5μM 9-*cis*-retinal (purple). The surface immunostaining intensities of HEK293T cells transiently expressing WT rhodopsin at 37°C (black) is shown for reference. C) Representative absorbance spectra of WT (black), G90M (red), and T289I (blue) rhodopsins that were regenerated in cellular membranes prior to purification are shown. D) Representative absorbance spectra of WT (black), G90M (red), and T289I (blue) rhodopsins that were regenerated in live cells prior to purification are shown.

## Discussion

Temperature-sensitive expression is an emergent property of class II mutations that serves as a marker for correctable disease variants within integral membrane proteins. However, many misfolded variants fail to recover their expression at lower temperatures or in the presence of pharmacological correctors. Basic insights into the molecular basis of these phenotypes are needed to rationalize the mutation-specific effects of corrector compounds.^32^ In this work, we employed DMS to measure the change in the PME of 700 rhodopsin variants at low temperature in relation to changes that occur in the presence of a stabilizing corrector molecule. Our findings suggest that, like WT, the PME of most rhodopsin variants is somewhat sensitive to temperature (Fig. 1). However, the coincidence between temperature- and retinal-sensitivity only emerges among mutations within certain regions of the protein (Figs. 2 & 3C). Based on computational estimates, we identify statistical trends suggesting deviations in the sensitivity of these variants stem from their combined influence on the energetics of cotranslational folding, the energetics of post-translational folding, and on the integrity of the retinal binding pocket (Table 1). Furthermore, an evaluation of purified temperature-sensitive variants suggests the proteostatic effects of retinal may often arise from its influence on cotranslational processes.

We have identified a cluster of temperature- and retinal-sensitive mutations within TM2 at residues that stabilize a native helical kink (Fig. 4). In previous investigations of bacteriorhodopsin (bR), Cao & Bowie found that a proline-mediated kink in helix B is stabilized by surrounding tertiary contacts that help to “pin” the helix into its native shape.^33^ This helix can be straightened by mutating this proline and deleting a flanking hydrogen bond.^33^ By comparison, we find the diglycine kink in TM2 is stabilized by a network of backbone and side chain hydrogen bonds (Fig. 4C). Mutations that either introduce non-native polar residues or delete native polar residues near this kink most often decrease PME^20^ and render rhodopsin sensitive to both temperature and retinal (Fig. 2 A & C). While bR retains considerable stability when helix B is straightened,^33^ our observations potentially suggest the stability of the kink in TM2 is critical for rhodopsin folding in the cell. In the case of G90D and T94I, the destabilization caused by the disruption of these hydrogen bonds (Fig. 4 D & E) is potentially offset by the formation of stabilizing non-native van der Waals^23^ or electrostatic^22^ interactions formed by the mutated side chains within the retinal binding pocket. Nevertheless, other surrounding RP mutants that alter these residues exhibit diminished PME and an enhanced sensitivity to temperature and retinal. Rationalizing the distinct phenotypes caused by these mutations will therefore require an understanding of how perturbations of this network differentially impact the photocycle and the stability of rhodopsin.

Our findings on TM7 build on our recent investigations showing that its polarity predisposes it to topological defects,^19^ and that the PME of rhodopsin is highly sensitive to the effects of mutations within this domain.^20^ In both previous investigations, we found that mutations that disrupt translocon-mediated membrane integration of the nascent chain are less sensitive to retinal,^19,20^ which could potentially arise from the kinetic constraints of topological isomerization relative to those of quality control.^1,34^ Similarly, we show here that TM7 variants are also less temperature-sensitive overall (Fig. 3B). However, our analysis across the entire mutational spectrum implies mutations that *enhance* membrane integration and/ or *stabilize* the native structure are among the least sensitive (Table 1, Fig. 5). Because TM7 contains several polar side chains involved in functional dynamics,^35,36^ including those of the NPXXY motif, most substitutions at these positions will increase its hydrophobicity. It therefore stands to reason that there may be an abundance of mutations in TM7 that can enhance its membrane integration. Likewise, many mutations within TM7 may stabilize the protein by suppressing the conformational dynamics that couple retinal isomerization to receptor activation.^35^ Accordingly, few mutations in TM7 (relative to TM2) are predicted to disrupt cotranslational membrane integration and/ or to further destabilize the native fold (Table 1). Mutations in TM7 are also more likely to perturb retinal binding, which is critical for the energetic coupling between binding and folding.^37^ Our data generally suggest mutations that are more distant from the binding pocket are more likely to be correctable. Taken together, the collective insensitivity of TM7 variants^20^ can be explained by a combination of factors that includes the attenuated response of destabilized variants as well as a heightened prevalence of mutations that stabilize the protein and/ or disrupt retinal binding.

Our biochemical characterizations of G90M and T289I rhodopsin highlight important caveats associated with underlying assumptions concerning the impacts of temperature and retinal on PME. While these mutations decrease the PME of the opsin apoprotein at 37°C, their expression exceeds that of WT at 27°C and at 37°C in the presence of retinal (Table 1). Such an expression profile could hypothetically emerge from an increase in the retinal binding affinity and in the stability of these mutant apoproteins at 27°C. However, our efforts to regenerate and purify these variants suggest that they bind retinal less efficiently than WT (Fig. 6C, Table 2) and have a lower kinetic stability at 27°C (Fig. S2). While we cannot rule out purification artifacts, these observations suggest the proteostatic response of these variants is unlikely to reflect an enhanced propensity to bind retinal and/ or to fold at lower temperatures. The fact that the regeneration of these variants is more efficient when retinal is added during translation in live cells suggests instead that these effects arise from differences in the fidelity of cotranslational folding and/ or quality control (Figs. 6 C & D). It should be noted that the regeneration of P23H rhodopsin, a variant that causes RP, is also more efficient when retinal analogs are added to living cells.^38^ Understanding the molecular basis of these observations may therefore help to inform efforts to develop efficacious correctors for retinopathies caused by rhodopsin misfolding.^16^

Together, our findings demonstrate how the molecular effects of mutations on the energetics of cotranslational folding, post-translational folding, and cofactor/ corrector binding can together produce diverse proteostatic and pharmacological profiles. Taking these mechanistic factors into consideration may therefore help overcome current challenges associated with the prediction of the effects of mutations and small molecules on protein stability and cellular proteostasis- an imminent challenge in precision medicine.^1,29,32^

## Materials & Methods

### Plasmid Preparation and Mutagenesis

PME measurements for individual rhodopsin variants were carried out using vectors produced via site directed mutagenesis on a previously described rhodopsin expression vector^18,19^ bearing and N-terminal hemagglutinin (HA) tag and a 3’ cassette containing an internal ribosome entry site (IRES) and an enhanced green fluorescent protein (eGFP) that marks transfected cells with eGFP. Vectors used to purify untagged rhodopsin variants were produced by cloning human rhodopsin into a pcDNA5.1 vector, and mutations were introduced by site directed mutagenesis. The preparation of mixed genetic libraries of TM2 and TM7 variants were described previously.^20^

### Deep Mutational Scanning

DMS data were generated using protocols described previously,^20^ except that intensity scores for individual missense variants were normalized relative to the average scores for the collection of nonsense variants within each helix. Normalized intensities measured at 27°C or in the presence of 5 μM 9-*cis*-retinal at 37°C were then normalized relative to that for each corresponding variant at 37°C in the absence of retinal. Intensity ratios represent the average value from two independent biological replicates. To restrict our analysis to high quality measurements, we omitted variants that failed to pass a series of previously described quality filters^20^ as well as those with intensity ratios that varied by more than a value of 0.4 across the two replicates.

### Surface Immunostaining of Transiently Expressed Rhodopsin Variants

To compare the effects of mutations, the PME of transiently expressed rhodopsin variants was assessed in HEK293T cells. Briefly, HEK293T cells were grow in Gibco Dulbecco’s Modified Eagle Medium (Life Technologies, Waltham, MA) supplemented with 10% fetal bovine serum and penicillin/ streptomycin at the indicated temperature in an incubator containing 5% CO_2_. Individual variants were transiently transfected using Lipofectamine 3000 (Invitrogen, Carlsbad, CA) two days prior to surface immunostaining with a Dylight550-conjugated anti-HA antibody (Invitrogen, Carlsbad, CA) and cellular fluorescence profiles were quantified by flow cytometry, as was previously described.^18,19^

### Regeneration and Purification of Rhodopsin Variants

Mutant rhodopsin pigments were expressed and purified as previously described.^18,19^ Briefly, HEK293T cells were transiently transfected with mutant expression vectors using polyethylenimine. Forty-eight hours post-transfection with cells from twenty 10-cm plates were washed with PBS, harvested, centrifuged at 800 g, and the pellet was either kept at −80°C or immediately resuspended in a buffer composed of 20 mM Bis-tris propane (BTP), 120 mM NaCl, and protease inhibitor cocktail (pH 7.5). 9-*cis*-retinal was added to the cell suspension from a DMSO stock solution to a final concentration of 10 μM, and then cells were incubated in the dark for 2 h at 4 °C on a nutator. The cell suspension was then lysed with n-dodecyl-β-D-maltopyranoside (DDM) added to a final concentration of 20 mM, followed by an incubation for 1 h at 4 °C on a nutator. The lysate was centrifuged at 100,000 g for 1 h at 4°C and rhodopsin pigments were purified from the supernatant by immunoaffinity chromatography using 200 μl of 1D4 anti-Rho antibody immobilized on CNBr-activated agarose (6 mg/ mL 1D4). Rhodopsin binding was performed for 1 h at 4°C. The resin was then transferred to a column and washed with 15 ml of buffer composed of 20 mM BTP, 120 mM NaCl, and 2 mM DDM (pH 7.5). Pigments were eluted with the same buffer, supplemented with 0.6 mg/ml of the TETSQVAPA peptide. UV-visible spectra were recorded from freshly purified rhodopsin samples in the dark using UV-visible spectrophotometer (Cary 60, Varian, Palo Alto, CA).

### Kinetic stability of purified rhodopsin

Purified rhodopsin samples were incubated at 27°C in the dark and their spectra were recorded every 2 minutes for 2 hours. Absorbance values at 485 nm were normalized relative to the value at the initial time point. Normalized intensities were then plotted as a function of time. All samples were measured in triplicate.

### Computational estimates, distance measurements, and statistical analyses

The estimated energetic effects of mutations on the free energy of folding were calculated using a membrane-protein optimized version of the Rosetta ΔΔG protocol,^28,39^ as was previously described.^19,20^ The estimated energetic effects of mutations on the energetics of translocon-mediated membrane integration were calculated using the ΔG Predictor web server (https://dgpred.cbr.su.se).^40^ The distance between the C_α_; of each amino acid and the centroid position of retinal was calculated based on the crystallographic structure of WT bovine rhodopsin (PDB 3C9L). Hydrogen bonding networks were identified after processing each crystallographic structure with Schrödinger’s Protein Preparation Wizard, which was used to build in missing protons (including those on histidine side chains) and optimize hydrogen bond assignments based on the local environment (Schrödinger Inc, New York, NY).^41^ Correlation coefficients were calculated, and statistical hypothesis testing was carried out using OriginLab 2019 software (OriginLab Corporation, Northampton, MA).

### Rigidity estimates based on the simulated thermal denaturation of rhodopsin

To evaluate the rigidity of the native contacts formed by the residues of TM2, we analyzed the 50 lowest energy models of a previously described homology model of human opsin^19^ using the constrained network analysis web server (https://cpclab.uni-duesseldorf.de/cna/main.php?#FOCUS).^26^ The resulting percolation and rigidity indices were taken as complementary measures of rigidity.^27^

## Acknowledgments

We thank Jeff Brodsky for helpful discussions. We thank Christiane Hassel and the Indiana University Flow Cytometry Core Facility for technical support. We thank the Indiana University Center for Genomics and Bioinformatics for experimental and bioinformatic support. This research was supported in part by grants from the National Institutes of Health (NIH) (R01GM129261 to J.P.S. and EY025214 to B.J.). F.J.R. acknowledges receipt of a predoctoral fellowship from the Graduate Training Program in Quantitative and Chemical Biology at Indiana University (T32 GM109825).

