## Supplemental Materials for "Systematic Profiling of Temperature- and Retinal-Sensitive Rhodopsin Variants by Deep Mutational Scanning"

[†]Authors contributed equally

*This File Includes:*
Figure S1
Figure S2

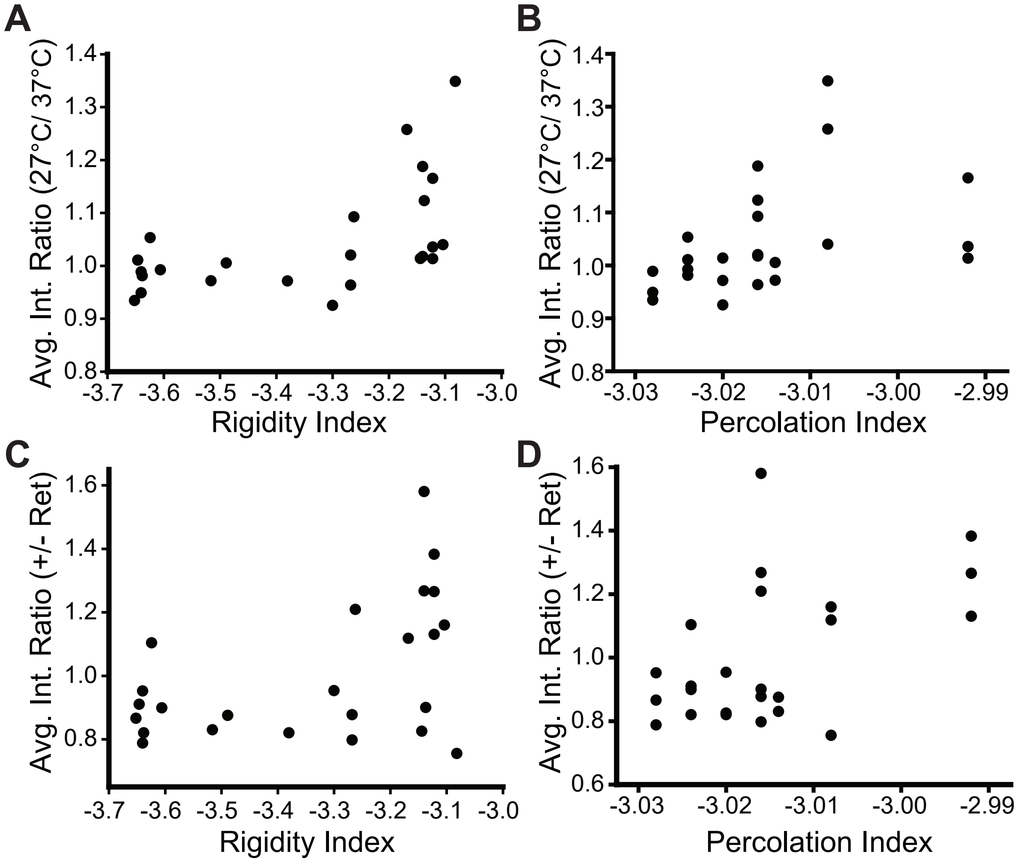


**Figure S1. Structural Rigidity of TM2 Residues in Relation to the Tempertat-ure- and Retinal- Sensitivity of TM2 Variants.** The ratio of the surface immu-nostaining intensity at 27°C versus 37°C was averaged accross all amino acid substitutions at each TM2 residue and plotted against the corresponding rigidity index (A) or percolation index (B) values that were derived for each residue from simulations of thermal unfolding. The ratio of the surface immunostaining intensi-ty in the presence and absence of 5 μM 9-*cis*-retinal was averaged accross all amino acid substitutions at each TM2 residue and plotted against the correspond-ing rigidity index (C) or percolation index (D) values that were derived for each residue from simulations of thermal unfolding. Lower rigidity and percolation index values, which are two measures of rigidity derived from simulations of ther-mal denaturation,[21] correspond to more structural rigidity. Percolation and rigidity index values ranged from -6 to 0, where a value of 0 corresponds to residue that completely lacks rigidity within the simulation and -6 corresponds to residues that retained rigidity the longest.

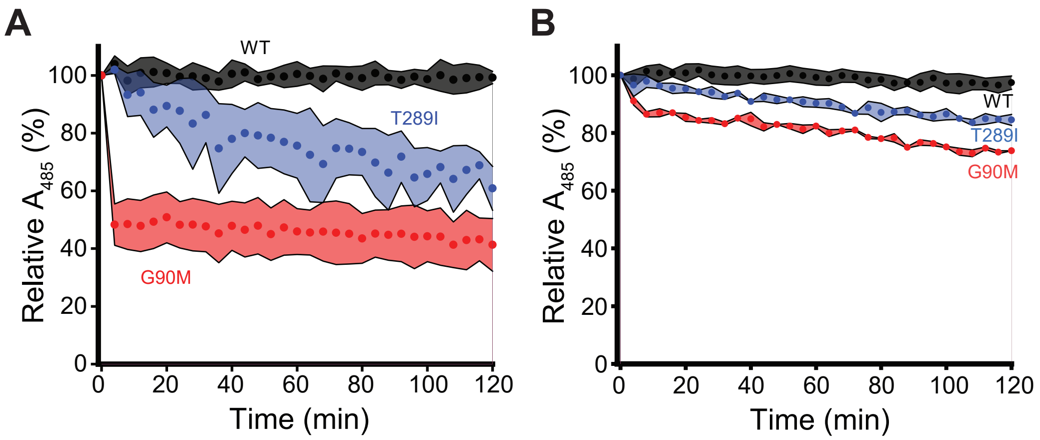


**Figure S2. Kinetic Stability of Purified G90M and T289I Rhodopsins.** A) WT (black), G90M (red), and T289I (blue) rhodopsins regenerated within cellular membranes prior to purification were incubated at 27°C, and the change in the absorbance at 485 nm was measured over time. Points reflect the average value and the shading reflects the bounds of the standard deviation from three experimental replicates. B) WT (black), G90M (red), and T289I (blue) rhodopsins regenerated within living cells prior to purification were incubated at 27°C, and the change in the absorbance at 485 nm was measured over time. Points reflect the average value and the shading reflects the bounds of the standard deviation from three experimental replicates.
